# Identification of a novel begomovirus betasatellite and occurrence of a viral complex associated with leaf curl disease in Bhut Jolokia in Assam

**DOI:** 10.1101/2024.11.12.623222

**Authors:** Richita Saikia, Ricky Raj Paswan, Basanta Kumar Borah

## Abstract

Bhut Jolokia (*Capsicum chinense* Jacq.) is one of the hottest chillies and an economically important crop in the northeastern regions of India. A major problem in the cultivation of Bhut Jolokia is the occurrence of leaf curl disease, which causes huge annual economic losses. The present study reveals the viral complex associated with the leaf curl disease in Bhut Jolokia collected from different geographical locations in Assam. A novel begomovirus betasatellite component was identified in two Bhut Jolokia samples, which was phylogenetically distinct from known begomovirus betasatellites. DAS-ELISA and PCR analysis confirmed the presence of multiple plant viruses including Chilli leaf curl virus (ChiLCV), Cucumber mosaic virus (CMV), Potato virus Y (PVY) and Tomato leaf curl virus (ToLCV) in most of the naturally infected Bhut Jolokia samples. Phylogenetic analysis revealed that the *coat protein* (*CP*) gene of ChiLCVs and ToLCVs were highly conserved compared to the *CP* of CMV.

## Introduction

The family, Solanaceae, comprises some of the most widely cultivated crops worldwide. Species belonging to this family, including pepper (*Capsicum* spp.), potato (*Solanum tuberosum*), tobacco (*Nicotiana tabacum*) and tomato (*S. lycopersicum*) are grown in both temperate and tropical climates (Hancinsky *et al*., 2020). *Capsicum chinense* Jacq. commonly known as Bhut Jolokia, or Ghost chilli is an indigenous solanaceous chilli variety cultivated in the northeastern region (NE) of India, mainly in Assam, Manipur, Meghalaya, Mizoram, and Nagaland (Chanu *et al*., 2017), and in other parts of the world (Liu and Nair, 2010). Bhut Jolokia is rich in vitamin A and C, and other essential nutrients, and consumed mainly for its pungency and flavour (Yogindran *et al*., 2021). The agro-climatic conditions of the NE states of India are favourable for the cultivation of Bhut Jolokia. Most *Capsicum* spp. contain capsaicin, an active ingredient responsible for their pungency. Bhut Jolokia, with a pungency exceeding 1 million Scoville Heat Units (SHUs), currently ranks 15th in heat level (The Hottest Peppers In The World, 2024 Update). Once recognized as the world’s hottest chili, it held the Guinness World Record in 2007 (Bosland and Baral, 2007). They possess anti-inflammatory and antioxidant properties and enhance the release of hydrolytic enzymes from saliva, which help in the digestion of starch (Surh, 2002). Bhut Jolokia occupies a crucial place in the different northeastern cuisines and is also exported as spice and condiments to other states of India and abroad.

The production of Bhut Jolokia is affected by the occurrence of leaf curl disease which causes huge economic losses every year (Talukdar *et al*., 2015). Leaf curl disease is a viral disease caused by a combination of multiple plant viruses which account for 60% of diseases that occurred in Bhut Jolokia in Assam (Talukdar *et al*., 2015). The symptoms include upward and downward leaf curling, mosaic patterns in the leaf, leaf deformation, leaf puckering, and bud necrosis (Meetei *et al*., 2020). Although multiple plant viruses belonging to the genus begomovirus, cucumovirus and potyvirus have been reported to be associated with the leaf curl disease in Bhut Jolokia (Banerjee *et al*., 2014; Baruah and Debnath, 2015; Talukdar *et al*., 2015; Meetei *et al*., 2020; Yogindran *et al*., 2021), the genetic diversity of these viral species infecting Bhut Jolokia has not been studied well. In the present study, we aimed to identify the viral complex and their associated betasatellite components in naturally infected Bhut Jolokia cultivated in different geographical locations in Assam, India. Here we identified and sequenced a conserved region of the *coat protein* gene from three different plant viruses, namely, ChiLCV, ToLCV, and CMV collected from infected Bhut Jolokia samples in addition to a novel ChiLCV betasatellite.

## Materials and method

### Field survey and sample collection

A systematic survey of Bhut Jolokia growing areas in Jorhat, Golaghat, and Sivasagar districts of Assam, India was conducted from February to April 2019 and 2021 (Fig. 1). Symptomatic leaves with leaf curling were collected from the Bhut Jolokia growing areas and the disease symptoms were recorded after that. The collected samples were stored at ‘-80°C’ until further use for nucleic acid extraction and Enzyme-linked immunosorbent assay (ELISA) test.

**Fig. 1.**
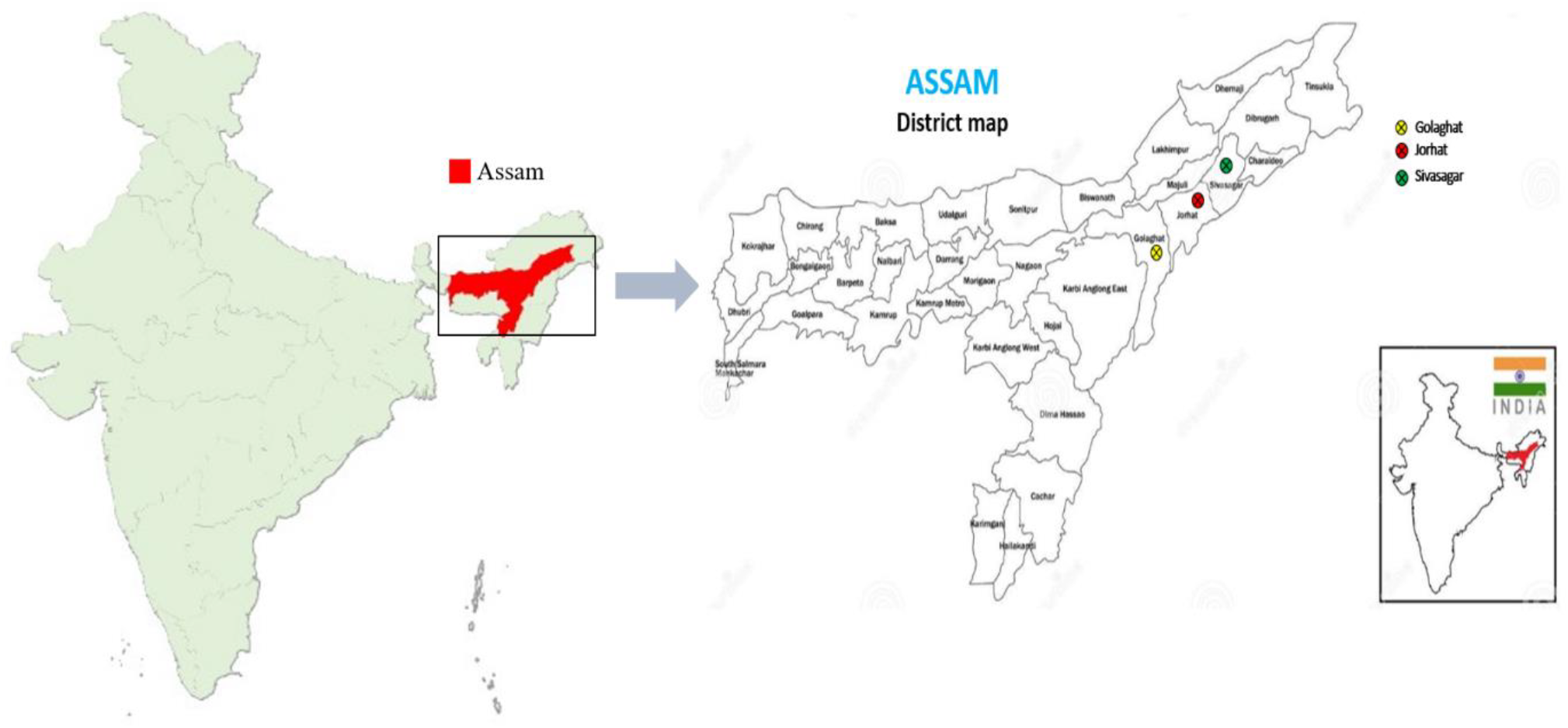
Collection of symptomatic leaf samples of Bhut Jolokia (*Capsicum chinense* Jacq.) from Assam, India. The yellow, red and green crossed circles represent samples collected from the Golaghat, Jorhat and Sivasagar districts of Assam, respectively.

### Virus detection and indexing

#### Serological detection

Double antibody sandwich enzyme-linked immunosorbent assay (DAS-ELISA) of symptomatic samples was carried out against three plant viruses, namely, CMV, PVY and TYLCV, along with their respective positive controls using the DAS-ELISA Kit (Bioreba AG, Switzerland) following the manufacturer’s instructions. The assay was performed on a 96-well polystyrene microtiter plate and the absorbance was recorded using the ELISA plate reader (BioRad, USA) at 405 nm wavelength. The values were considered positive if the absorbance was two times greater or equal to the negative control, as per standard protocol.

### Nucleic acid extraction and PCR amplification

#### Detection of DNA viruses in Bhut Jolokia

To detect candidate DNA viruses, total genomic DNA was isolated from symptomatic leaves using the Cetyl-trimethylammonium bromide (CTAB) method (Doyle and Doyle, 1990). To validate the DAS-ELISA results, PCR was performed using the coat protein gene-specific primers (Supplementary Table S2). A total of 20 μL reaction volume was used containing 1 µL of template DNA, 2 µL of 10X Advance PCR buffer (Promega, USA), 2 µL of 10 mM dNTPs, 0.5 µL of 10 µM each forward and reverse primers, 1 µL of *Pfu* DNA polymerase (2-3 U/µL, Promega, USA) and the remaining volume was adjusted with nuclease free water. PCR amplification was performed at 94°C for 3 min for initial denaturation, which was followed by 35 cycles of 94°C for 45 sec, primers annealing at 52°C for ChiLCV and 46°C for TYLCV for 30 sec, and 72°C for 1 min with a final extension at 72°C for 7 min. PCR for detecting beta satellites was carried out following the same PCR profile except annealing at 60°C for 30 sec. A geminivirus universal betasatellite primers (Briddon *et al*., 2002) were used to detect and identify novel viral sequences in the Bhut Jolokia samples. The details of the primers used are provided in Supplementary Table S2.

#### Detection of RNA viruses in Bhut Jolokia

Total RNA was extracted from symptomatic Bhut Jolokia leaf samples using the TRI Reagent® (Sigma, USA). The quantity and integrity of the isolated RNA were checked using the NanoDrop™ 1000 spectrophotometer (Thermo Scientific, USA) and agarose gel electrophoresis. cDNA was synthesized from the total RNA using PrimeScript™ 1st strand cDNA Synthesis Kit (Takara, Japan). A 20 μL of reaction volume was used containing 1 μg of total RNA template, 4 μL of 5X PrimeScript buffer, 1 μL of 50 μM oligo dT primers, 1 µL of 10 mM dNTPs, 0.5 µL of 40 U/µL RNase Inhibitor, 0.5 µL of PrimeScript RTase (200 U/µL) and the remaining volume was adjusted with nuclease-free water. The cDNA synthesis was performed in a thermocycler at 42°C for 60 min followed by 95°C for 5 min denaturation. For the reverse transcription (RT) PCR, 1 µL of template cDNA was used in a total reaction volume of 20 µL, which includes 2 µL of 10X Advance PCR buffer (Promega, USA), 2 µL of 10 mM dNTPs, 0.5 µl of 10 µM each forward and reverse primer, 1 µl of *Pfu* DNA polymerase (2-3 U/µl, Promega, USA) and 13.5 µl nuclease free water. PCR was performed at 94°C for 3 min for initial denaturation, followed by 35 cycles of 94°C for 45 sec, primer annealing at 58°C for CMV *CP* 50°C for ChiVMV *CP* and 56.5°C for PVY *CP* for 30 sec, 72°C for 1 min and with a final extension at 72°C for 7 min. The amplified RT-PCR products were run in 1.5% ethidium bromide-stained agarose gel electrophoresis at 100 V for 45 min. The PCR products were gel purified and sequenced by Sanger sequencing (Eurofins Genomics, Karnataka, India).

#### Transmission assay

The infection assay involved mechanically inoculating healthy plants by applying sap from naturally infected plants. Infected plant material (1g) was taken to grind using a mortar and pestle in 0.01 M phosphate buffer (pH 7.0) and the homogenate was filtered through a muslin cloth. The test plants that were used were young and tender, dusted with carborundum powder. Approximately, 60 µl sap was applied to the systemic leaves by gently wiping onto each marked leaf. The inoculated leaves were washed after some time with water. The inoculated plants (*Amaranthus caudatus, Nicotiana benthamiana*, and *Capsicum chinense* Jacq.) were then transferred to a growth cabinet. After symptom development, PCR with gene-specific primers was used to confirm viral presence.

#### Sequence analysis

The sequences obtained from Sanger sequencing were compared with other publicly available viral sequences available in the NCBI GenBank database (https://www.ncbi.nlm.nih.gov/) using the BLASTn (Basic Local Alignment Search) tool. For phylogenetic analysis, the MUSCLE alignment tool was used to align the sequences of the closely related taxa followed by the analysis of sequence divergence by the Molecular Evolutionary Genetic Analysis (MEGA11). The phylogenetic relatedness of the sequences obtained was analysed using the maximum likelihood method (ML) with 1000 bootstrap replicate values.

Based on the per cent nucleotide identity, a color-coded matrix (heat map) of the pairwise similarity scores were generated using the sequence demarcation tool (SDT, version 1.2) for the sequences of the cognate viruses and betasatellites (Muhire *et al*., 2014). The heat map provided a comprehensive and visual representation of the genetic relationships and similarities among the viral and betasatellite sequences under investigation.

## Results

### Field survey and collection of samples

Bhut Jolokia plants showing symptoms of upward and downward leaf curling, leaf puckering, vein banding and leaf deformation, mosaic mottling, and stunted growth were collected from Golaghat, Jorhat and Sivasagar districts of Assam (Fig. 1, 2, Table S1). A total of 29 symptomatic samples were collected from Golaghat (4 samples), Jorhat (23 samples), and Sivasagar (2 samples). The distribution shows that most symptomatic plants occurred in Jorhat, implying either higher susceptibility of Bhut Jolokia to viral infections or environmental conditions that promote disease spread. Whitefly infestation was commonly observed in the Bhut jolokia plants with leaf curl symptoms (Fig. 2f). The presence of virus was confirmed by serological and nucleic acid-based detection assays. Mechanical transfection of infected Bhut Jolokia samples was carried out on three different host plants, namely Amaranthus, Tobacco and Bhut Jolokia to check for viral transmissibility and the host spectrum of the disease complex (Fig. 3 and Supplementary Fig. 1)

**Fig. 2.**
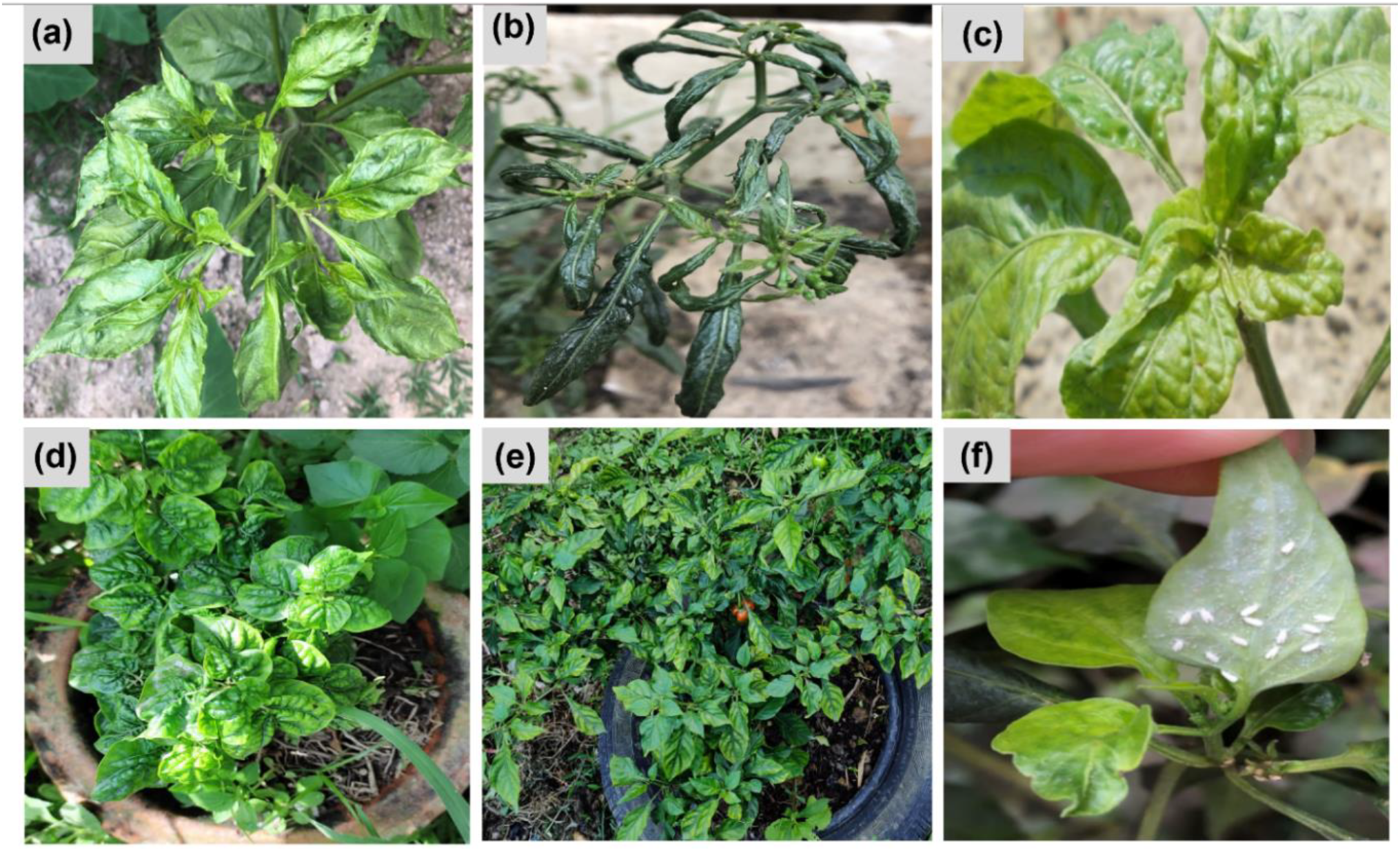
**(a-e)** Visible leaf curl disease symptoms observed in the Bhut Jolokia samples collected from different geographical locations in Assam. (a) leaf narrowing. (b) Downward curling. (c) leaf puckering. (d) vein banding and leaf deformation. (e) mosaic mottling and stunted growth (f) whitefly infestation.

**Fig. 3.**
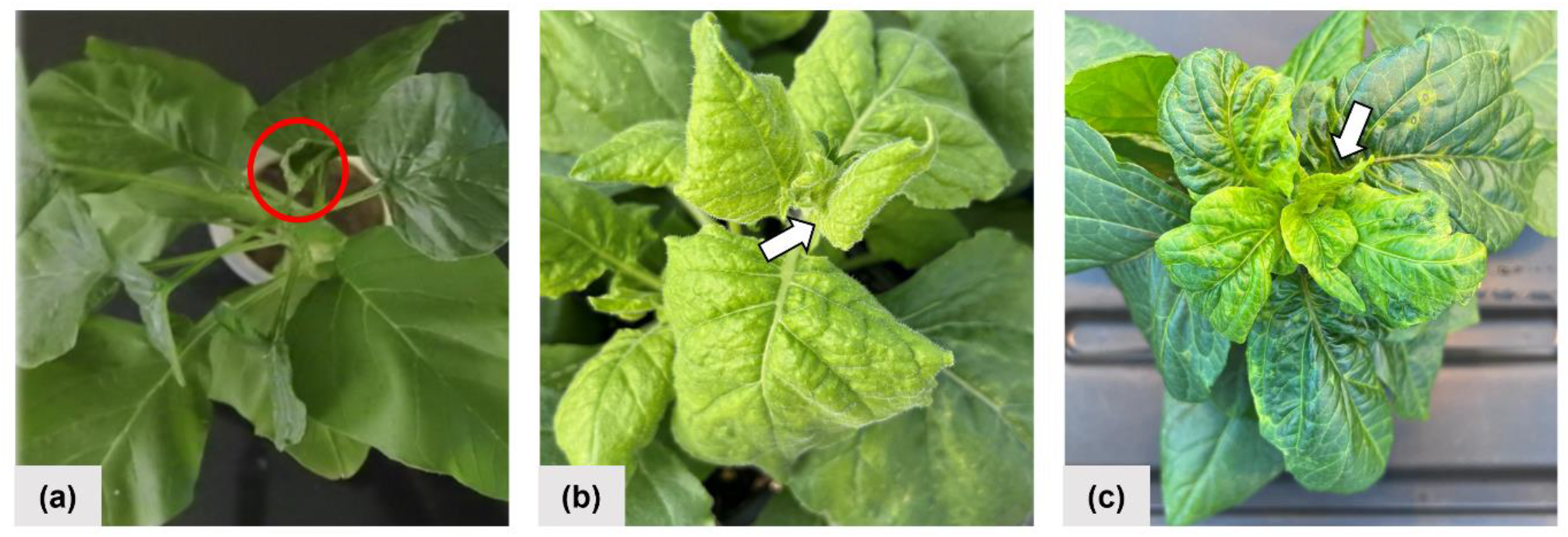
Virus transmissibility and development of leaf curl disease in three host plants that were inoculated mechanically with saps from symptomatic Bhut Jolokia samples, (a) *Amaranthus caudatus*. (b) *Nicotiana benthamiana* (c) *Capsicum chinense* Jacq. Photos were taken three weeks after sap inoculation from symptomatic Bhut Jolokia leaves. Red circle and white colour arrow indicate leaf curl symptom in different host plants.

### Serological and molecular detection of plant viruses

DAS-ELISA assay detected the presence of three plant viruses, namely ChiLCV, CMV and PVY belonging to the genus begomovirus, cucumovirus and potyvirus, respectively. All the twenty-three naturally infected Bhut Jolokia samples from Jorhat district were found to be positive for ChiLCV, CMV and PVY (Table 1). The samples collected from the Golaghat and Sivasagar districts were not positive for any of these viral groups. DAS-ELISA positive samples from Jorhat were further subjected to molecular screening through PCR.

**Table 1.** Summary of different plant viruses detected in symptomatic Bhut Jolokia samples collected from Jorhat district.

| Sample ID<sup>a</sup> | Viruses detected |  |  |  |  |  |  |
| --- | --- | --- | --- | --- | --- | --- | --- |
|  | ChiLCV | ToLCV | ToLCNDV | Beta-sat | CMV | PVY<sup>b</sup> | ChiVMV |
| JH1 | + | + | - | - | + | + | - |
| JH2 | - | + | - | - | - | + | - |
| JH3 | - | - | - | - | + | + | - |
| JH4 | + | + | - | - | - | + | - |
| JH5 | - | + | - | - | - | + | - |
| JH6 | - | - | - | - | + | + | - |
| JH7 | - | - | - | - | + | + | - |
| JH8 | - | - | - | - | - | - | - |
| JH9 | - | - | - | - | - | - | - |
| JH10 | - | - | - | - | - | - | - |
| JH11 | - | - | - | - | - | - | - |
| JH12 | - | - | - | - | - | - | - |
| JH13 | - | + | - | - | + | - | - |
| JH14 | - | + | - | - | - | - | - |
| JH15 | - | - | - | - | - | - | - |
| JH16 | + | + | - | - | + | - | - |
| JH17 | + | + | - | - | - | - | - |
| JH18 | - | - | - | - | + | - | + |
| JH19 | + | + | - | + | + | - | - |
| JH20 | - | + | - | - | - | - | - |
| JH21 | + | + | - | + | + | - | - |
| JH22 | - | + | - | - | - | - | - |
| JH23 | - | - | - | - | - | - | - |

<sup>a</sup>The plus (+) sign indicates PCR and DAS-ELISA positive samples for the respective plant viruses, whereas the minus (-) sign indicates samples that were not positive.
<sup>b</sup>All PVY positive (+) samples were based on only DAS-ELISA results.

PCR analysis of the Bhut Jolokia samples targeting the conserved coat protein (*CP*) genes of the respective plant viruses revealed mixed viral infections. Out of the total symptomatic Bhut Jolokia samples, six samples were PCR positive for ChiLCV (*CP*, 443 bp) showing 20.7% incidence rate, 13 samples were positive for ToLCV (*CP*, 578 bp) resulting in 44.8% incidence rate and nine samples tested positive for CMV (*CP*, 454 bp), giving 31% incidence rate (Table 1 and Fig. S2). These results suggest that ToLCV was the most prevalent virus among the tested symptomatic samples, followed by CMV and ChiLCV. Although all the samples collected from Jorhat district were positive for PVY as determined by DAS-ELISA, none exhibited positive results when subjected to PCR analysis (Table 1). All the samples, except one (JH18) tested negative for both Tomato leaf curl New Delhi virus (ToLCNDV) and Chilli veinal mottle virus (ChiVMV). Mixed infection of at least three different plant viruses, *viz*., ChiLCV, ToLCV and CMV, were observed in four samples: JH1, JH16, JH19 and JH21 (Table 1); suggesting possible association of several plant viruses with leaf curl disease in Bhut Jolokia. Similarly, two of the infected Bhut Jolokia samples (JH19 and JH21) from Jorhat revealed the presence of betasatellite (amplicon size of ∼1.35 Kb in length) (Fig. 4).

**Fig. 4.**
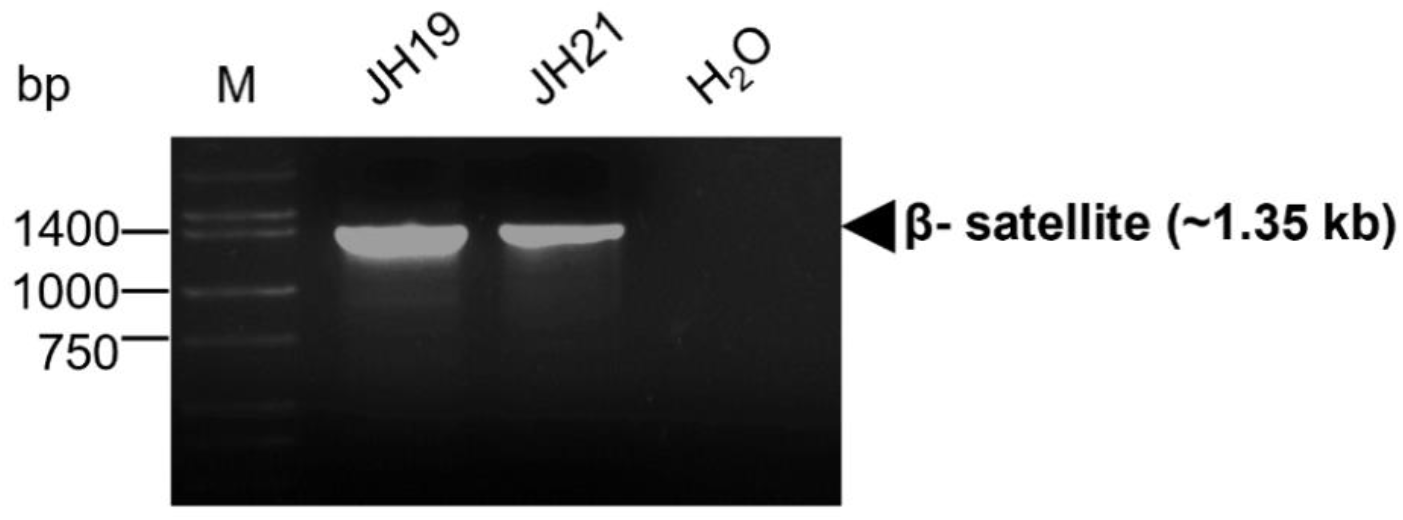
Detection of a betasatellite associated with the leaf curl disease in Bhut Jolokia. Full length PCR amplification of betasatellite components using the universal begomovirus specific betasatellite primers from Briddon *et al*., 2002. JH19 and JH21 represent Bhut Jolokia samples with leaf curl disease collected from two locations from Jorhat, Assam. The arrowhead on the right indicates the amplified target gene fragments. H2O indicate template-free negative control. M indicates 50-10,000 bp DNA ladder (Takara, Japan).

### Sequencing of viral amplicons and sequence analyses

The contigs derived from the sequenced PCR amplicons were checked and deposited in the NCBI database. The submitted sequences along with their GenBank accession numbers and sequence lengths are listed in Table 2. The viral sequences were annotated based on their highest nucleotide identity. The BLASTn results confirmed the presence of ChiLCV, CMV, and ToLCV, detected by DAS-ELISA and PCR analysis in the infected Bhut Jolokia samples collected from Jorhat district. The sequencing of betasatellite amplicons from two Bhut Jolokia samples (JH19 and JH21) was similarly validated through BLASTn analysis. The sequences isolated from infected Bhut Jolokia samples with leaf curl symptoms for ChiLCV had 97.47% (JH1), 97.25% (JH16) and 95.73% (JH21) identity with the Pepper leaf curl Bangladesh virus-India [India/Ghazipur/2007] (Accession no. HM007097.1) and 96.58% (JH19) identical with the Chilli leaf curl Cooch Behar (Accession no. MN851261.1) (Table 3). Similarly, ToLCV had 98.36% (JH1), 97.69% (JH16), 97.51% (JH19) and 97.86% (JH21) identity to the Tomato leaf curl Joydebpur virus clone (Accession no. JQ654461.1) (Table 3). In case of CMV, sequence JH1 exhibited 98.02% identity to the CMV isolate from pepper plants found in Guangdong and Shandong, China (Accession no. FJ403474.1). Moreover, sequences JH16 and JH21 showed similarities of 99.09% and 98.80% identity (Table 3), respectively, to the CMV isolate from watermelon in Yunnan, China (Accession no. OP617565.1). Additionally, the sequence of JH19 displayed 98.31% identity (Table 3) to the CMV isolate from hot pepper plants in Thailand (Accession no. AY560556.1).

**Table 2.**
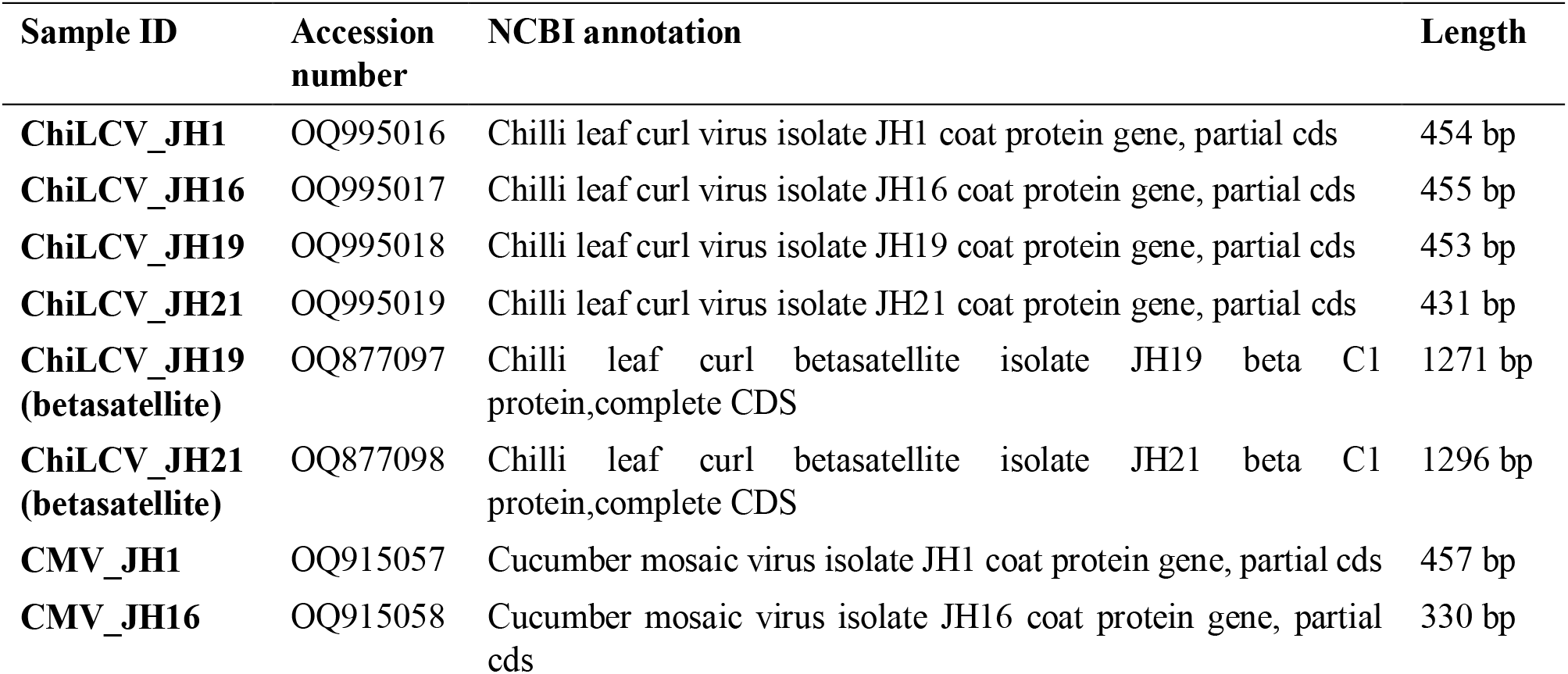
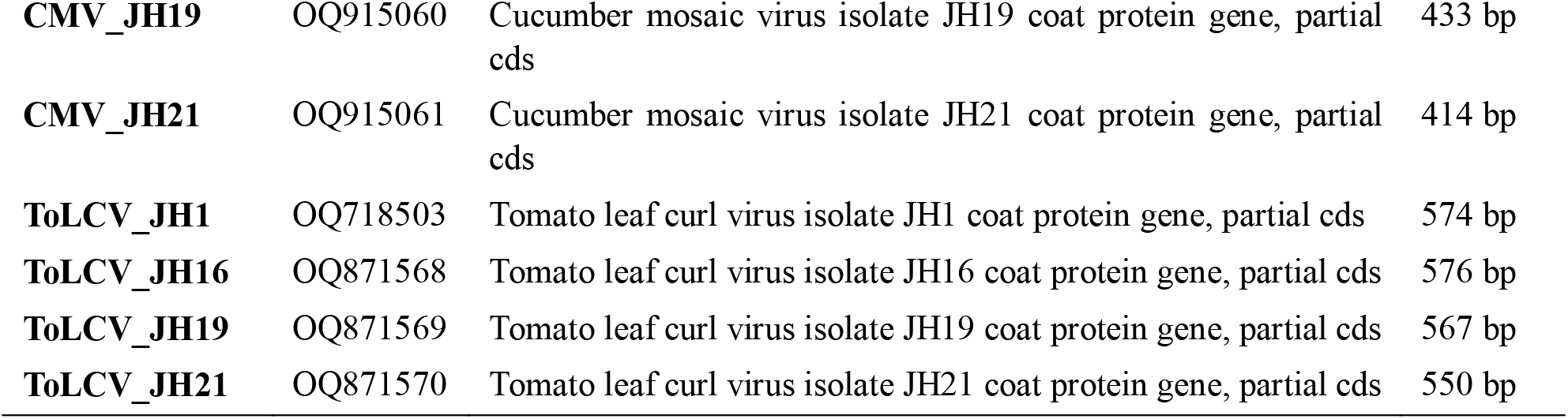
List of sequences of the coat protein genes of ChiLCV, CMV, ToLCV, as well as the betasatellite deposited obtained from Bhut Jolokia samples, deposited in the GenBank.

**Table 3.**
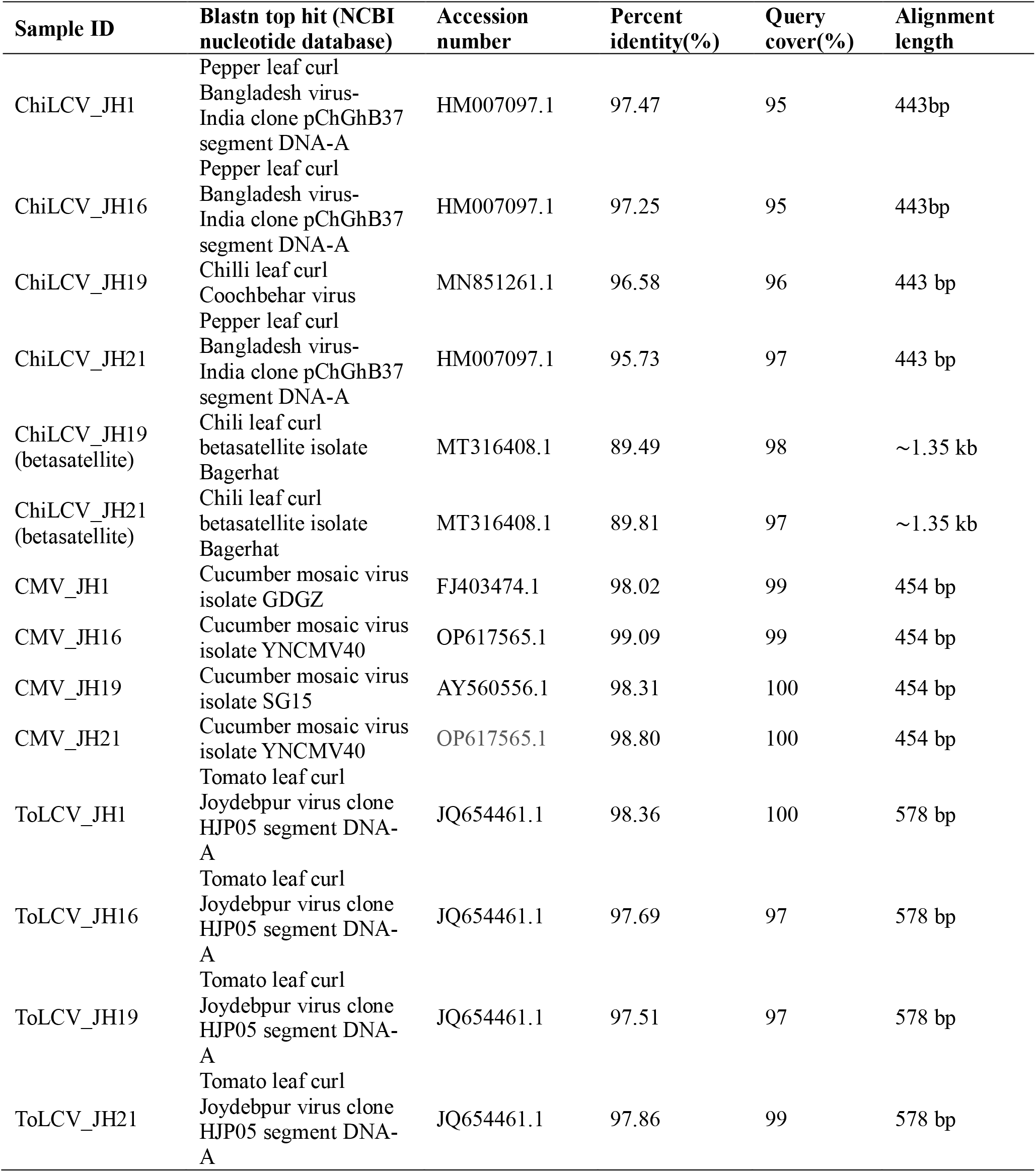
The top-most BLASTn hits of the *coat protein* gene sequences of ChiLCV, CMV, ToLCV and the associated betasatellite sequences with the publicly available viral sequences.

The sequencing of beta-satellite amplicons, which is approximately 1.35 kb, covered the entire genome size except for the primer binding site (Fig. S3). The sequencing of beta-satellite amplicons from the two Bhut Jolokia samples (JH19 and JH21) gave two full-length sequences of varied lengths 1271 bp and 1296 bp, respectively (Table 2). The betasatellite samples, JH19 and JH21 exhibited identities of 89.49% and 89.81%, respectively, to the betasatellite isolate Bagerhat from chili pepper plants in Bangladesh (Accession no. MT316408.1) (Table 3). The nucleotide similarity-search results for all the obtained partial and full-length sequences are listed in Table 3. The detected sequences from each virus isolate along with other publicly available viral isolates, multiple sequence alignment was conducted followed by phylogenetic tree construction (Fig. 5 and Fig.6).

**Fig. 5.**
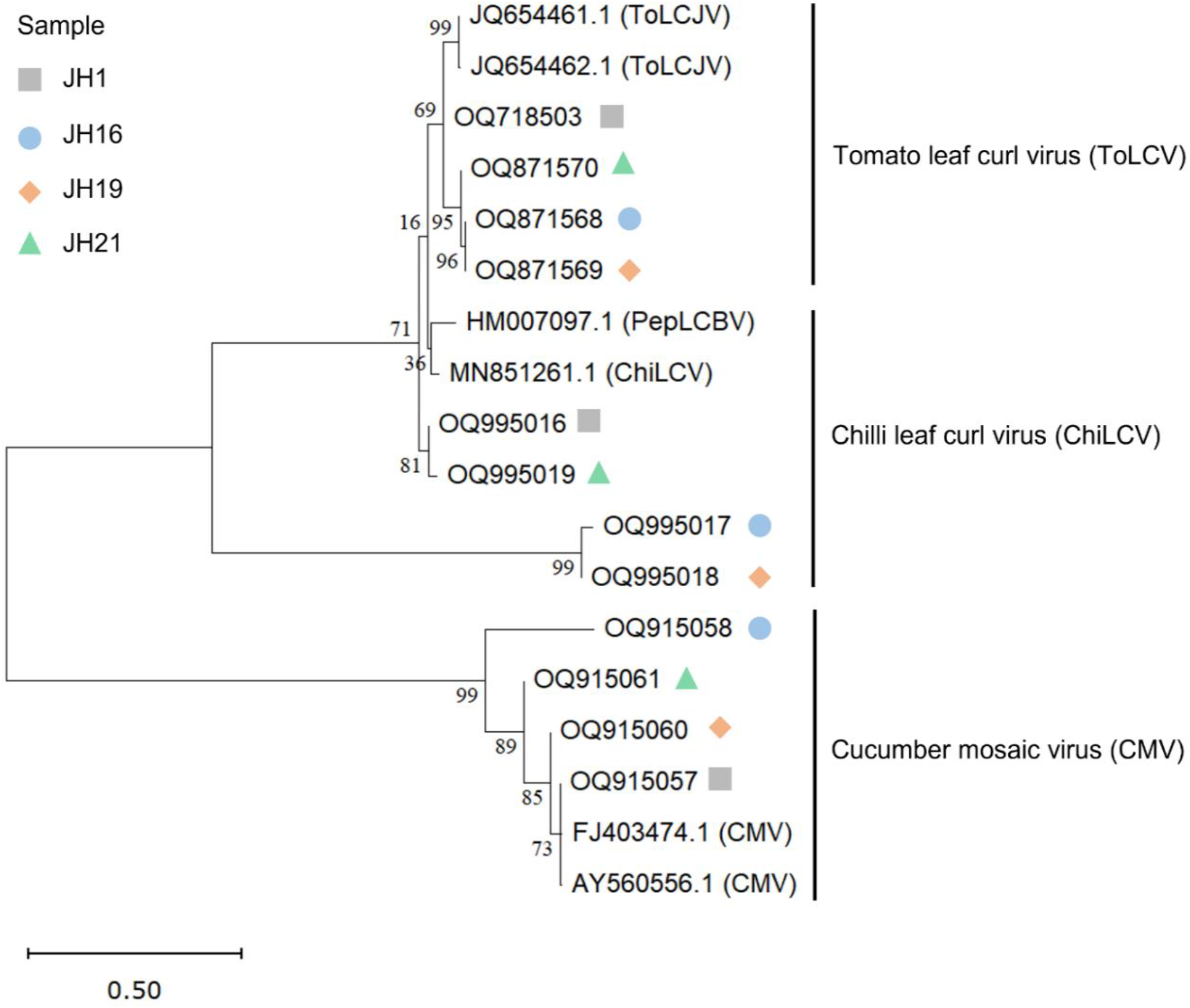
Phylogenetic relatedness based on the *coat protein (CP)* gene sequences of Tomato leaf curl virus (ToLCV), Chilli leaf curl virus (ChiLCV) and Cucumber mosaic virus (CMV) obtained from selected Bhut Jolokia samples with mixed viral infections (Table 4.6). Publicly available coat protein gene sequences of the corresponding plant viruses (ToLCV, ChiLCV and CMV) were collected and a phylogenetic tree was constructed with the Maximum likelihood method with 1000 bootstrap replicates using MEGAv11. The CMV clade was used as an outgroup for ToLCV and ChiLCV and vice versa.

### Phylogenetic analysis

To understand the relationship of the obtained viral sequences and the betasatellite sequences from the four samples, i e, JH1, JH16, JH19 and JH21, with other publicly available viral isolates, phylogenetic trees were constructed using the maximum-likelihood method. The sequences related to ChiLCV formed a coherent cluster with each other and with the sequences of ToLCV as shown in the phylogenetic tree (Fig. 6). It was observed that all the four ToLCV samples (JH1, JH16, JH19 and JH21) were closely related to each other, while JH16, JH19 and JH21 formed a coherent cluster with each other by bootstrap confidence of 95% and 96%, thereby suggesting their genetic relatedness. While all four ToLCV samples formed a coherent cluster with JQ654461.1 (Tomato leaf curl Joydebpur virus clone HJP05 segment DNA-A) and JQ654462.1 (Tomato leaf curl Joydebpur virus clone HJP07 segment DNA-A) with a bootstrap confidence of 69%. All ToLCV samples (JH1, JH16, JH19 and JH21) and two ChiLCV samples (JH1 and JH21) formed a clade with the previously reported isolates of ToLCV from Mungbean (accessions: JQ654461.1 and JQ654462.1) and ChiLCV from Chilli (accessions: HM007097.1 and MN851261.1). This observation implies the presence of genetic similarities between ChiLCV and ToLCV, which maybe possibly due to shared evolutionary history or common genetic features. The two ChiLCV isolates, JH16 and JH19, formed a distinct clade, which is clearly separated from the rest of the ChiLCV and ToLCV isolates with a bootstrap confidence of 71%. On the other hand, all four CMV isolates sequenced in the present study, JH1, JH16, JH19, JH21 (accessions: OQ915057, OQ915058, OQ915060, OQ915061, respectively) formed a cluster with each other and with the previously reported CMV isolates reported from hot pepper, indicating their genetic relatedness (accessions: FJ403474.1, Cucumber mosaic virus isolate GDGZ coat protein (CP) gene and AY560556.1, Cucumber mosaic virus isolate SG15 coat protein gene). As expected, all the sequences obtained from the symptomatic Bhut Jolokia samples collected from Jorhat corresponding to the *CP* gene of ChiLCV, CMV and ToLCV (JH1, JH16, JH19 and JH21) clustered well into three different viral groups. Each viral family acted as an outgroup to each other indicating their distinct evolutionary origins (Fig. 6). The associated betasatellite sequences obtained from the infected Bhut Jolokia samples, JH19 and JH21 (accession no. OQ877097 and OQ877098, respectively) formed a distinct clade in the phylogenetic tree, which segregated from the remaining publicly accessible betasatellite sequences (Fig. 7). The betasatellite samples, JH19 and JH21 exhibited a considerable degree of similarity, based on bootstrap confidence of 100%.

**Fig. 6.**
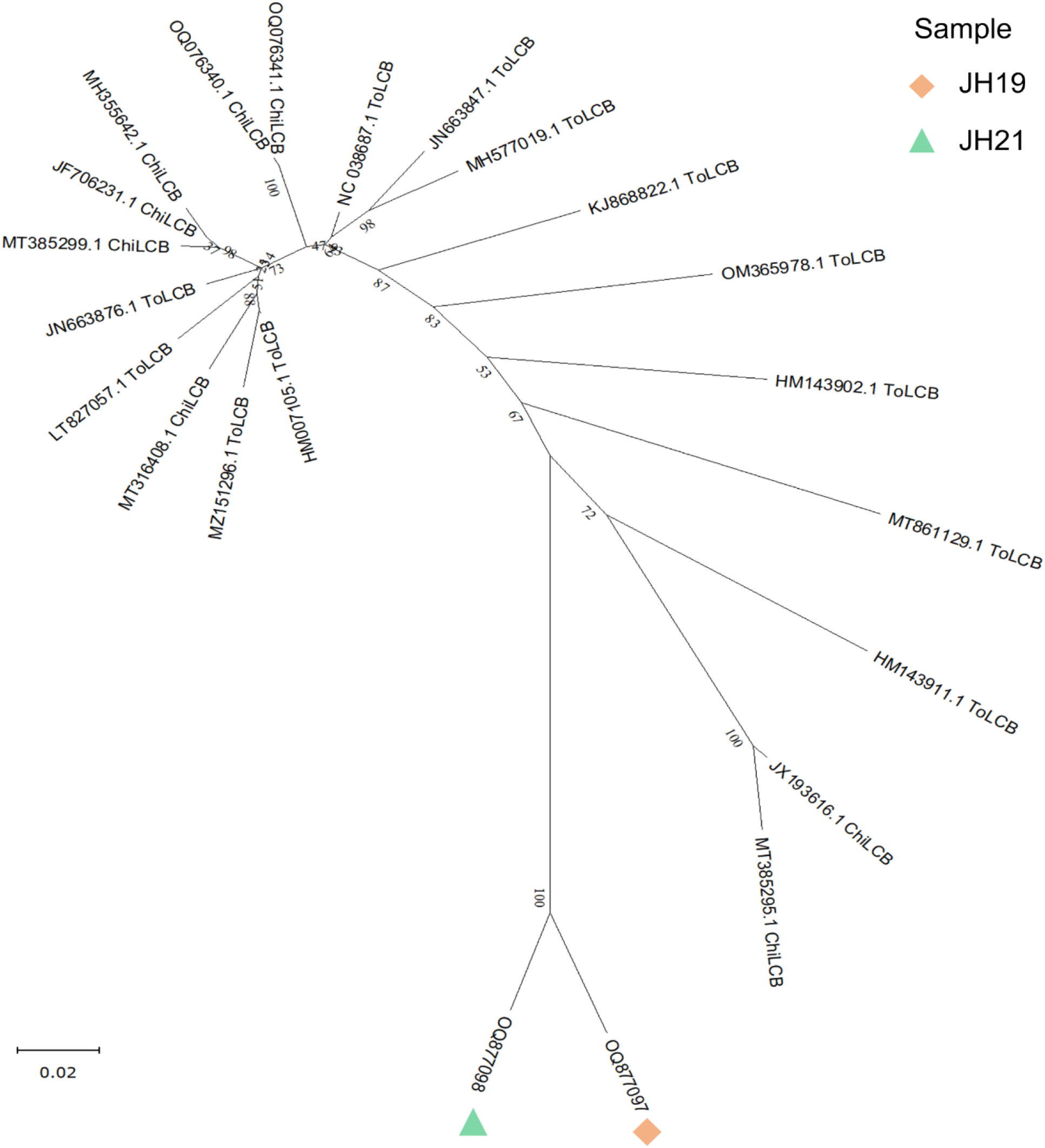
Phylogenetic relatedness based on the betasatellite sequences obtained from the naturally infected Bhut Jolokia samples, JH19 and JH21, and other publicly available betasatellite sequences from the NCBI nucleotide database. The Maximum likelihood method was used to construct the phylogenetic tree with 1000 bootstrap replicates using MEGA v11.

**Fig. 7.**
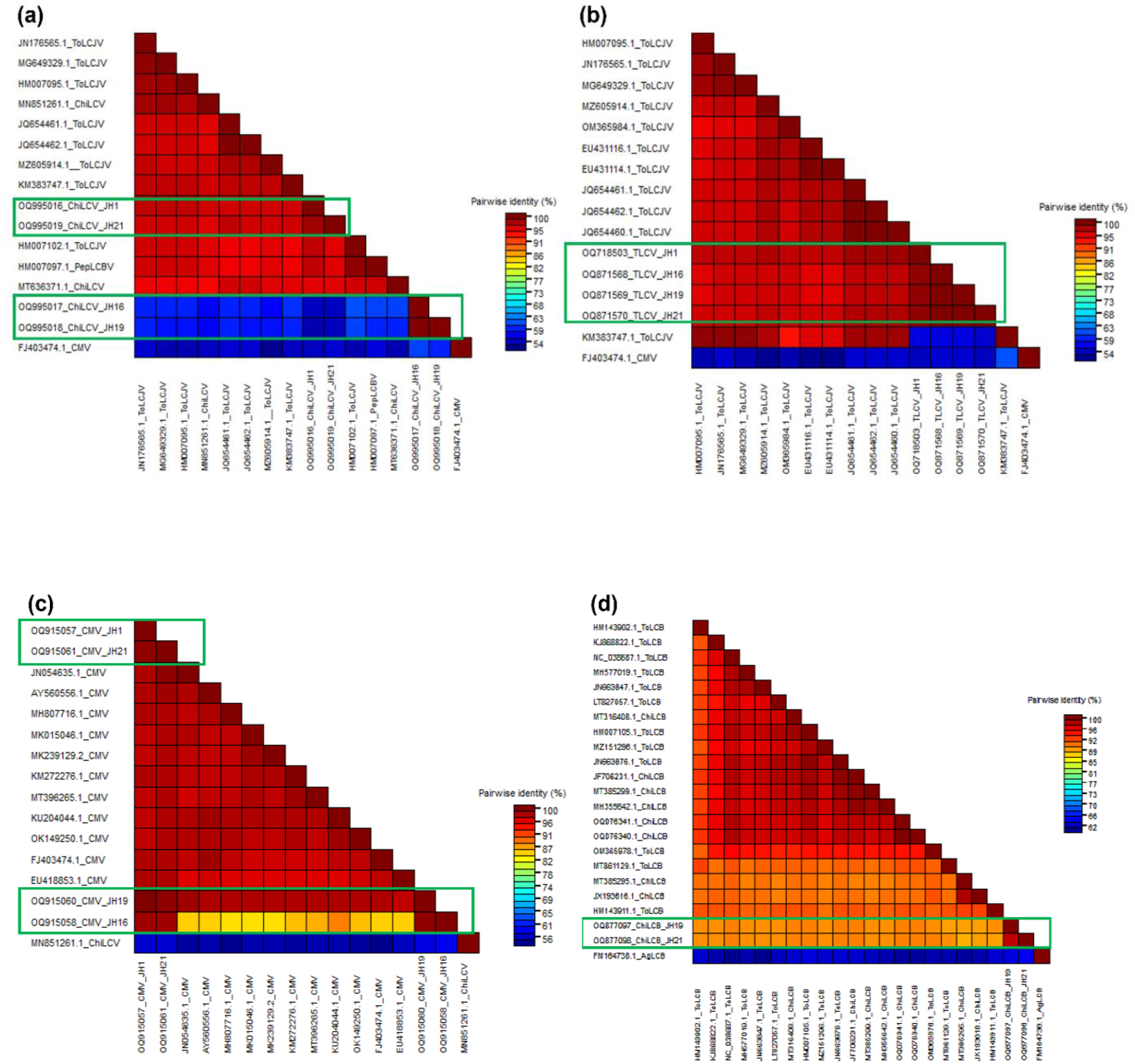
Percent nucleotide identity-based heat map based on *coat protein* (*CP*) gene created using sequence demarcation tool (SDT), version 1.2. for (a) Chilli leaf curl virus. The *CP* gene of Cucumber mosaic virus (GenBank accession no.FJ403474.1) was considered as an outgroup for the analysis (b) Tomato leaf curl virus. The *CP* gene Cucumber mosaic virus (GenBank accession no.FJ403474.1) was considered as an outgroup for the analysis (c) Cucumber mosaic virus. The *CP* gene of Chilli leaf curl virus (GenBank accession no. MN851261.1) Chilli leaf curl virus betasatellites. The Ageratum leaf curl disease associated betasatellite (GenBank accession no. FM164738.1) was considered as an outgroup for the analysis.

### Percent nucleotide identity-based heat map

A color-coded matrix (heat map) of pairwise similarity scores was generated using SDT v1.2 for the sequences of the cognate viruses and betasatellites based on the per cent nucleotide identity. The heat map provided a comprehensive and visual representation of the genetic relationships and similarities among the viral and betasatellite sequences under investigation which further confirmed the results obtained from the phylogenetic analysis (Fig. 8).

The ChiLCV *CP* sequences from JH16 and JH19 samples exhibit significant genetic divergence (per cent identity < 70%) compared to JH1, JH21, and other publicly available ChiLCV sequences (Fig. 7a). The ToLCV samples (JH1, JH16, JH19, JH21) exhibit a considerable genetic similarity (>90%) with other ToLCV strains reported in different geographical regions of India (Fig. 7b). The CMV sequence from JH16 displays significant genetic divergence compared to other CMV sequences from remaining Jorhat samples and other CMV samples in general, suggesting a distinct lineage or strain specific to JH16 (Fig. 7c). The two betasatellite samples, JH19 and JH21, identified as Chilli leaf curl virus betasatellites, exhibited less than 90% pairwise identity (Fig. 7d) with other publicly available betasatellite sequences.

## Discussion

*Capsicum chinense* Jacq., commonly known as Bhut Jolokia, is an indigenous chili cultivar native to North-east India. An important challenge in the cultivation of *C. chinense* Jacq. is the frequent incidence of leaf curl disease, leading to substantial crop yield losses. Despite a few previous reports about the disease and the causal pathogens, a comprehensive understanding of the disease, the pathogens and the manifestation of the disease symptoms remains limited. The present study reveals the presence of multiple plant viruses in the infected Bhut Jolokia samples, indicating that the leaf curl disease is attributed to a viral complex rather than a single virus along with the association of satellite molecules. Several plant viruses have been previously reported in Bhut Jolokia from north-east region of India including Assam (Banerjee *et al*., 2014; Talukdar *et al*., 2015, 2017; Baruah *et al*., 2016; Meetei *et al*., 2020 and Yogindran *et al*., 2021). However, leaf curling and similar symptoms are challenging to interpret because they closely mimic the effects of abiotic stresses, especially in the initial phases of nitrogen deficiency in plants (Jeong et al., 2014).

Mixed virus infections play a crucial role in virus evolution as they create opportunities for recombination, potentially giving rise to more virulent strains or novel begomovirus species (Kumar *et al*., 2019). There are reports of occurrence of at least five viruses associated with the leaf curl disease of Bhut Jolokia. They are: CMV, PVY, GBNV, ChiLCV and TSWV (Talukdar *et al*., 2015), ChiVMV (Banerjee *et al*., 2014; Meetei *et al*., 2020) and CLCuMuV (Yogindran *et al*., 2021). According to the findings of Baruah *et al*. (2016), Bhut Jolokia exhibited viral infection with multiple plant viruses, leading to the formation of a disease complex. Similar findings have been observed in other chilli cultivars (Khan *et al*., 2006; Wang and Bosland, 2006 and Meena *et al*., 2008). The etiology of leaf curl disease in chilli or pepper has been ascribed to the association with more than twenty distinct species of geminiviruses (Yogindran *et al*., 2021). Furthermore, the clustering patterns within the dendrograms using the MEGA-11 provide valuable insights into the genetic diversity and evolution of the studied viruses. ToLCV and ChiLCV samples formed a clade, suggesting genetic similarities, possibly due to shared evolutionary history or common genetic features (Fig. 5). The two detected betasatellites from the infected Bhut Jolokia samples from Jorhat are novel betasatellite sequences and were therefore named Chilli leaf curl betasatellite based on the similarity with other Chilli leaf curl betasatellites in the NCBI database (Fig. 6). The nucleotide sequences of ChiLCV showed low per cent identity indicating that they might belong to distinct subtypes or even different species within the begomoviral group. These findings suggest the presence of a novel ChiLCV strain which might be responsible for leaf curl disease in Bhut Jolokia in the region of Assam. Other known ChiLCV strains have been detected in different regions of India and Bangladesh (Venkataravanappa *et al*., 2016; Kumar *et al*., 2015), suggesting that co-existence of multiple ChiLCV strains is possible in the northeastern part of India. The *CP* gene of the ToLCVs detected in the Bhut Jolokia samples showed high sequence similarities with each other and with other previously reported Tomato leaf curl Joydebpur virus (ToLCJV) isolates (Fig. 7b). The result suggests that leaf curl disease of Bhut Jolokia is caused by a combination of multiple plant viruses viz. ChiLCV, ToLCV and CMV. The occurrence of these viral groups in Bhut Jolokia might occurred as an event of host jump from their original hosts (such as chilli and tomato). Further studies are needed to understand the evolutionary history of this viral group and host selection mechanisms.

The International Committee on Taxonomy of Viruses (ICTV) based on the proposal by Brown *et al*., (2015) set as a demarcation threshold of 91% and 94% pairwise nucleotide identity for considering new begomovirus species and strains, respectively (Brown *et al*., 2015). In the present study, the two betasatellite components had a pairwise nucleotide identity of less than 90%, suggesting a novel betasatellite component detected in Bhut Jolokia from Jorhat, Assam. Blastn search of the novel betasatellites showed close resemblance with the betasatellites from known ChiLCV isolates and the result correlates well with the pairwise nucleotide alignment performed in the present study (Fig. 7d). The above results suggest that the novel betasatellites (accessions: OQ877097 and OQ877098) identified in the present study is possibly associated with the ChiLCV detected samples JH19 (accession: OQ995018) and JH21 (accession: OQ995019). The identified betasatellites were possibly linked to the unique ChiLCV isolates detected in JH19 and JH21 based on the degree of similarity with other Chilli leaf curl betasatellites deposited in NCBI. Additionally, a high level of similarity between the two betasatellite samples in terms of per cent identity indicates that they could be variants of the same viral species. Chattopadhyay *et al*., (2008) reported the association of betasatellites with ChiLCV in leaf curl infected chili samples. Betasatellites play an additive role in the process of inducing disease symptoms, determining the host range, and overcoming host defense mechanisms (Briddon and Stanley, 2006). The occurrence of mixed infections involving various begomoviruses, satellite molecules and the emergence of novel virus variants through recombination of existing ones (Fiallo-Olivé *et al*., 2012) demonstrates their ability to adapt to new hosts, posing a significant threat to economically important crops (Varma and Malathi, 2003). Majority of the betasatellites are observed in association with Old World (OW) monopartite begomoviruses, while only a few are found to be linked with bipartite begomoviruses (Ashwathappa *et al*., 2022; Shingote *et al*., 2022). ChiLCV has a monopartite genome and belongs to the OW begomoviruses (Shingote *et al*., 2022). The identification of a novel betasatellite further suggests the involvement of a potentially novel strain of a virus, possibly related to Chilli leaf curl virus (Fig 6 and Fig 7d). The precise role of this betasatellite in enhancing the virulence of the associated virus remains uncertain. Therefore, additional complementation assays are needed, both with and without the cognate virus, in the host plant. Further comprehensive investigations are required to determine the viral complex of leaf curl disease in Bhut Jolokia. The outcome of this study could be valuable in the formulation and prioritization of virus management initiatives, encompassing the development of cultivars resistant to the virus, aimed at mitigating the impact of these viral infections within the specified region. Therefore, a thorough investigation is essential for the detection of other viruses, across all the regions where Bhut Jolokia is cultivated, and employing advanced methodologies for comprehensive coverage.

## Author contribution

This work was carried out in collaboration among all authors. RS and RRP performed the research. RS did the data analysis and wrote the first draft of the manuscript with inputs from all the authors. BKB conceived and designed the research, along with supervising the experiments.

## Funding

This work was funded by a DBT-sponsored project (Grant no-BT/PR25287/NER/95/1113/2017 (DBT-NER)), implemented in 2018.

## Acknowledgements

RS acknowledges the University Grants Commission (UGC), India for providing her with a National Fellowship and Scholarship for Higher Education (Award No - 202021-NFST-ASS-00455) to pursue her Ph.D. degree.

## Notes

### Competing Interest Statement

The authors have declared no competing interest.

