## Supplementary Materials for "Identification of a novel begomovirus betasatellite and occurrence of a viral complex associated with leaf curl disease in Bhut Jolokia in Assam"

**Supplementary files**

**Figure legends**

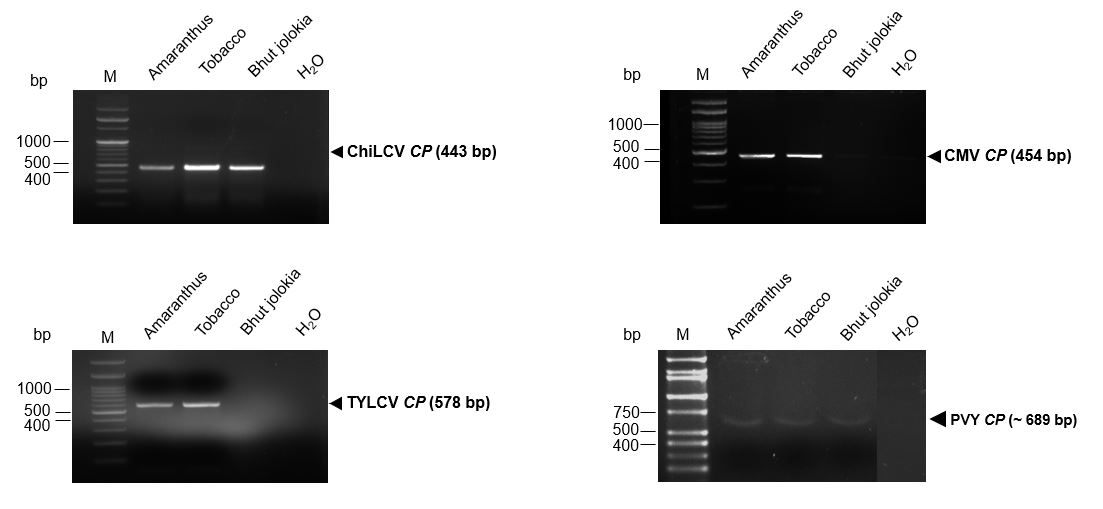

**Fig.1** Confirmation of viral components in mechanically inoculated host plants by PCR analysis. The *coat protein* (*CP*) genes of the cognate viruses were amplified using gene-specific primers and the amplified products were run in agarose gel electrophoresis (130V for 30 minutes). (a) *Chilli leaf curl virus* (ChiLCV *CP*, 443 bp). (b) *Tomato leaf curl virus* (ToLCV *CP*, 578 bp). (c) *Cucumber mosaic virus* (CMV *CP*, 454 bp). (d) *Potato virus Y* (PVY *CP*, 689 bp). Black colour arrows indicate the amplified target gene fragments. H_2_O indicates template-free negative control. M indicates 50-10,000 bp DNA ladder (Takara, Japan).

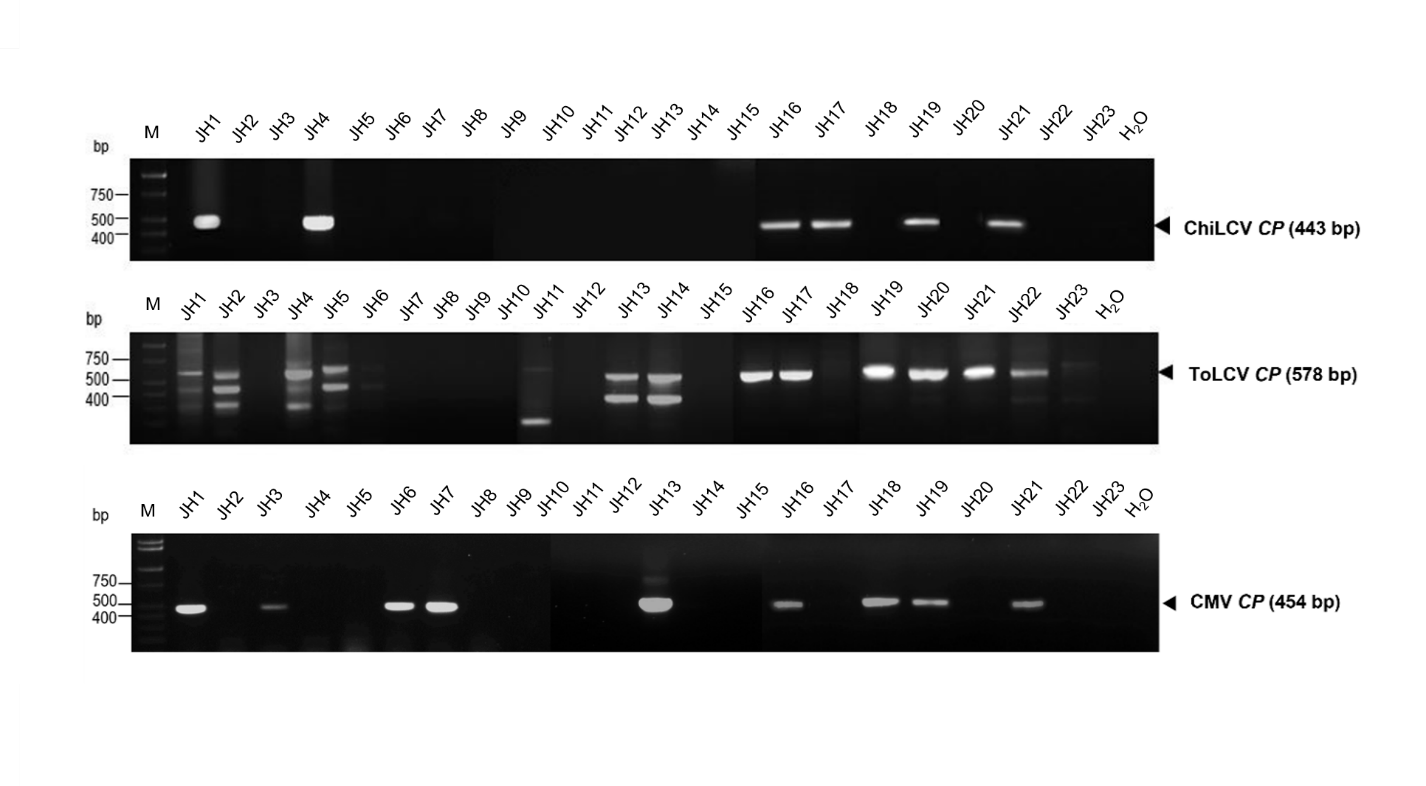

**Fig. 2** Detection of multiple plant viruses in symptomatic Bhut Jolokia samples collected from Jorhat district of Assam. The *CP* genes of the cognate viruses, viz., *Chili leaf curl virus* (ChiLCV), *Tomato leaf curl virus* (ToLCV) and *Cucumber mosaic virus* (CMV) were amplified by PCR using gene-specific primers. JH1-JH23 represents symptomatic Bhut Jolokia samples. The black arrowheads on the right indicate the amplified target gene fragments. H_2_O indicates template-free negative control. M indicates 50-10,000 bp DNA ladder (Takara, Japan).

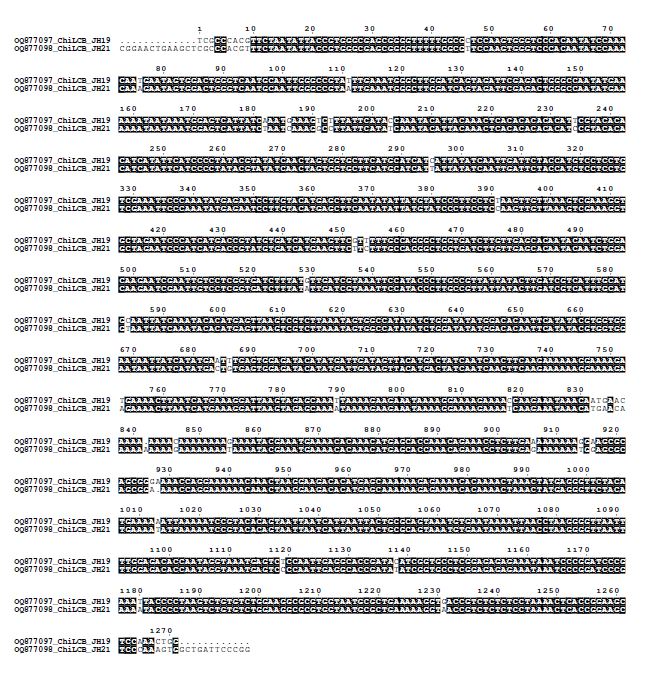

**Fig. 3** Sequence alignment of the two novel betasatellites (OQ877097_JH19 and OQ877098_JH21) associated with the leaf curl disease in Bhut Jolokia from Assam.

**Table S1. Symptomatic Bhut Jolokia samples collected from different geographical locations in Assam, India**

| SampleID | Symptoms observed in the host | Location | Coordinates (Latitude and Longitude) |
| --- | --- | --- | --- |
| JH1 | Upward curling of leaves and stunted growth | Nagabaat, Jorhat | 26°21'40.9"N 94°09'41.9"E |
| GH1 | Crinkling, puckering of the leaves, stunted growth. | Tengani Gaon, Golaghat | 26°24'22.2"N 93°55'56.6"E |
| GH2 | Mosaic and puckering of the leaves, stunted growth | Ahom Gaon, Golaghat | 26°27'58.7"N 94°01'14.4"E |
| GH3 | Upward curling and puckering of leaves, stunted growth | Tamuli Gaon, Golaghat | 26°54'39.4"N 94°28'32.5"E |
| JH2 | Downward curling, vein thickening and stunted growth | Horticultural Orchard, AAU, Jorhat | 26°43'38.7"N 94°12'05.3"E |
| JH3 | Puckering of the leaves and stunted growth | KVK, Kaliapani, Teok | 26°49'53.1"N 94°27'26.9"E |
| SG1 | Vein banding and puckering of the leaves, stunted growth | Notun Lunpuria Gaon, Namti, Sivasagar | 26°56'45.3"N 94°47'39.7"E |
| SG2 | Mosaic symptoms, stunted growth, and vein thickening | Dulakharia Gaon, Sivasagar | 26°53'42.9"N 94°27'44.1"E |
| GH4 | Stunted growth, vein clearing and small leaves | Jamuguri Gaon, Golaghat | 26°18'54.4"N 93°57'16.9"E |
| JH4 | Upward and downward curling of the leaves | Horticultural Orchard, AAU, Jorhat | 26°43'38.7"N 94°12'05.3"E |
| JH5 | Leaf puckering, mosaic symptoms and small leaves | Horticultural Orchard, AAU, Jorhat | 26°43'38.7"N 94°12'05.3"E |
| JH6 | Upward curling of leaves and small leaves | Horticultural Orchard, AAU, Jorhat | 26°43'38.7"N 94°12'05.3"E |
| JH7 | Leaf puckering, vein thickening and mosaic symptoms | Horticultural Orchard, AAU, Jorhat | 26°43'38.7"N 94°12'05.3"E |
| JH8 | Upward curling of leaves, puckering of leaves and stunted growth | AAU, Jorhat | 26°43'22.9"N 94°11'48.0"E |
| JH9 | Puckering of leaves, vein thickening and stunted growth | Barbheta, Jorhat | 26°43'30.6"N 94°11'51.9"E |
| JH10 | Mosaic symptoms, leaf puckering and stunted growth | Barbheta, Jorhat | 26°43'30.6"N 94°11'51.9"E |
| JH11 | Leaf puckering, vein banding and stunted growth | Barbheta, Jorhat | 26°43'30.6"N 94°11'51.9"E |
| JH12 | Mosaic symptoms, small leaves and crinkling | Barbheta, Jorhat | 26°43'34.3"N 94°11'46.4"E |
| JH13 | Leaf puckering and stunted growth | Nowchaliah Gaon, Jorhat | 26°43'34.0"N 94°11'28.0"E |
| JH14 | Mosaic and puckering of the leaves and stunted growth | Nowchaliah Gaon, Jorhat | 26°43'34.0"N 94°11'28.0"E |
| JH15 | Leaf puckering, vein banding and stunted growth | Barbheta, Jorhat | 26°43'31.5"N 94°11'51.9"E |
| JH16 | Upward curling of leaves, small leaves and stunted growth | Barbheta, Jorhat | 26°44'19.5"N 94°12'10.2"E |
| JH17 | Mosaic symptoms, crinkling and small leaves | Barbheta, Jorhat | 26°44'19.5"N 94°12'10.2"E |
| JH18 | Leaf puckering, mosaic symptoms and stunted growth | Barbheta, Jorhat | 26°44'19.5"N 94°12'10.2"E |
| JH19 | Upward curling of the leaves, vein thickening, small leaves | Rajabari, Jorhat | 26°44'29.7"N 94°13'54.0"E |
| JH20 | Downward curling, small leaves and stunted growth | Nowchaliah gaon, Jorhat | 26°43'34.0"N 94°11'28.0"E |
| JH21 | Downward curling of leaves, vein thickening, small leaves | Rajabari, Jorhat | 26°44'29.7"N 94°13'54.0"E |
| JH22 | Leaf puckering, stunted growth, and small leaves | Rajabari, Jorhat | 26°44'29.7"N 94°13'54.0"E |
| JH23 | Upward curling of the leaves and small leaves | Rowriah, Jorhat | 26°43'58.5"N 94°11'07.0"E |

**Supplementary Table S2**. **List of primers used for detection of the plant viruses in *C. chinense* Jacq.**

| **Primer name** | **Viral target** | **Sequence (5' to 3')** | **Amplicon size (bp)** | **Reference, if any** |
| --- | --- | --- | --- | --- |
| B01-F B02-R | Geminivirus satellite (Betasatellite) | GGTACCACTACGCTACGCAGCAGCC  GGTACCTACCCTCCCAGGGGTACAC | 1350 | Briddon *et al*.*,* 2002 |
| ChiLCV_AV1-F ChiLCV_AV1-R | *Chilli leaf curl virus* | GAAGCCCAGGATGTACAGGA CTAATGCCTGCTCCTTCGAC | 443 | - |
| ChiVMV_F ChiVMV_R | *Chilli veinal mottle virus* | TCTTGATTATGCCCCTGAGC  TCCTGTGTGCCTACCCTACC | 519 | - |
| CMV_AV1-F CMV_AV1-R | *Cucumber mosaic virus* | AACCAGTGCTGGTCGTAACC  TGCGGCATACTGATAAACCA | 454 | - |
| PVY_S5585M PVY_6266R | *Potato virus Y* | GGATCTCAAGTTGAAGGGGAC CTCCTGTGCTGGTATGTCCT | 689 | Lorenzen *et al*., 2006 |
| ToLCNDV-F ToLCNDV-R | Tomato leaf curl New Delhi virus | GAACTATGGTGAAGCGACCAGCAGA ACACAGGTCCTTAGGTACCTGG | 914 | Gawande *et al*.,2007 |
| TYLCV_1F TYLCV_1R | *Tomato yellow leaf curl virus* | GCCCATGTAYCGRAAGCC GGRTTAGARGCATGMGTAC | 578 | Accotto *et al*., 2000 |
